# CRISPRO Identifies Functional Protein Coding Sequences Based on Genome Editing Dense Mutagenesis

**DOI:** 10.1101/326504

**Authors:** Vivien A. C. Schoonenberg, Mitchel A. Cole, Qiuming Yao, Claudio Macias-Treviño, Falak Sher, Patrick G. Schupp, Matthew C. Canver, Takahiro Maeda, Luca Pinello, Daniel E. Bauer

## Abstract

CRISPR/Cas9 pooled screening permits parallel evaluation of comprehensive guide RNA libraries to systematically perturb protein coding sequences in situ and correlate with functional readouts. For the analysis and visualization of the resulting datasets we have developed CRISPRO, a computational pipeline that maps functional scores associated with guide RNAs to genome, transcript, and protein coordinates and structure. No available tool has similar functionality. The ensuing genotype-phenotype linear and 3D maps raise hypotheses about structure-function relationships at discrete protein regions. Machine learning based on CRISPRO features improves prediction of guide RNA efficacy. The CRISPRO tool is freely available at gitlab.com/bauerlab/crispro.

## Background

Clustered regularly interspaced short palindromic repeats (CRISPR) - Cas9 genome editing technologies permit new approaches for the dissection of gene function. Cas9 cleavage results in imprecise end-joining repair products with indels. Biallelic frameshift mutations lead to loss-of-function of the gene product, often through nonsense mediated decay (NMD) destabilizing the transcript. This paradigm allows for the systematic dissection of genetic dependencies in genome-wide CRISPR screens in the context of disease-relevant cellular phenotypes [1–3]. However, the mechanisms by which individual alleles contribute to cellular phenotypes are not typically directly assessed in these experiments. Such information could aid in the rational design of novel therapeutics as well as in the context of biological engineering to reprogram gene circuitry.

Following a programmable nuclease mediated double-strand break, the major genome editing outcome is imprecise end-joining, as produced by classical NHEJ and microhomology-mediated end joining pathways. The ensuing indel spectrum is comprised of short indels, typically up to 10-20 base pair (bp) in length. Although the distribution of indel length is non-uniform and depends on target sequence and cellular repair contexts, on average 2/3 of alleles from the indel spectrum of end-joining repair following an induced double strand break (DSB) result in frameshifts. For a gene with two genomic copies and independently assorting repair alleles, on average ˜4/9 of edited cells would be expected to produce a biallelic frameshift, causing complete loss-of-function. The remaining ˜5/9 of cells would retain partial gene function from in-frame alleles, assuming gain or loss of a short stretch of amino acids would be tolerated by the protein. However, guide RNAs targeting the coding sequence of critical residues may be associated with heightened functional impact within a population of cells by causing loss-of-function not only from frameshift but also from in-frame mutations [4]. Here we explore comprehensive dense mutagenesis with many cleavages per gene to systematically define functional protein coding sequences. This method is also known as a CRISPR tiling or guide RNA saturating mutagenesis screen. A typical design would be to include as many guide RNAs as possible, as restricted by a given protospacer adjacent motif (PAM) availability for a given nuclease (such as the NGG motif in the case of SpCas9) [5,6]. A single pooled screen experiment may employ large numbers of guide RNAs to systematically disrupt the function of numerous protein coding genes (Figure 1A).

**Figure 1.**
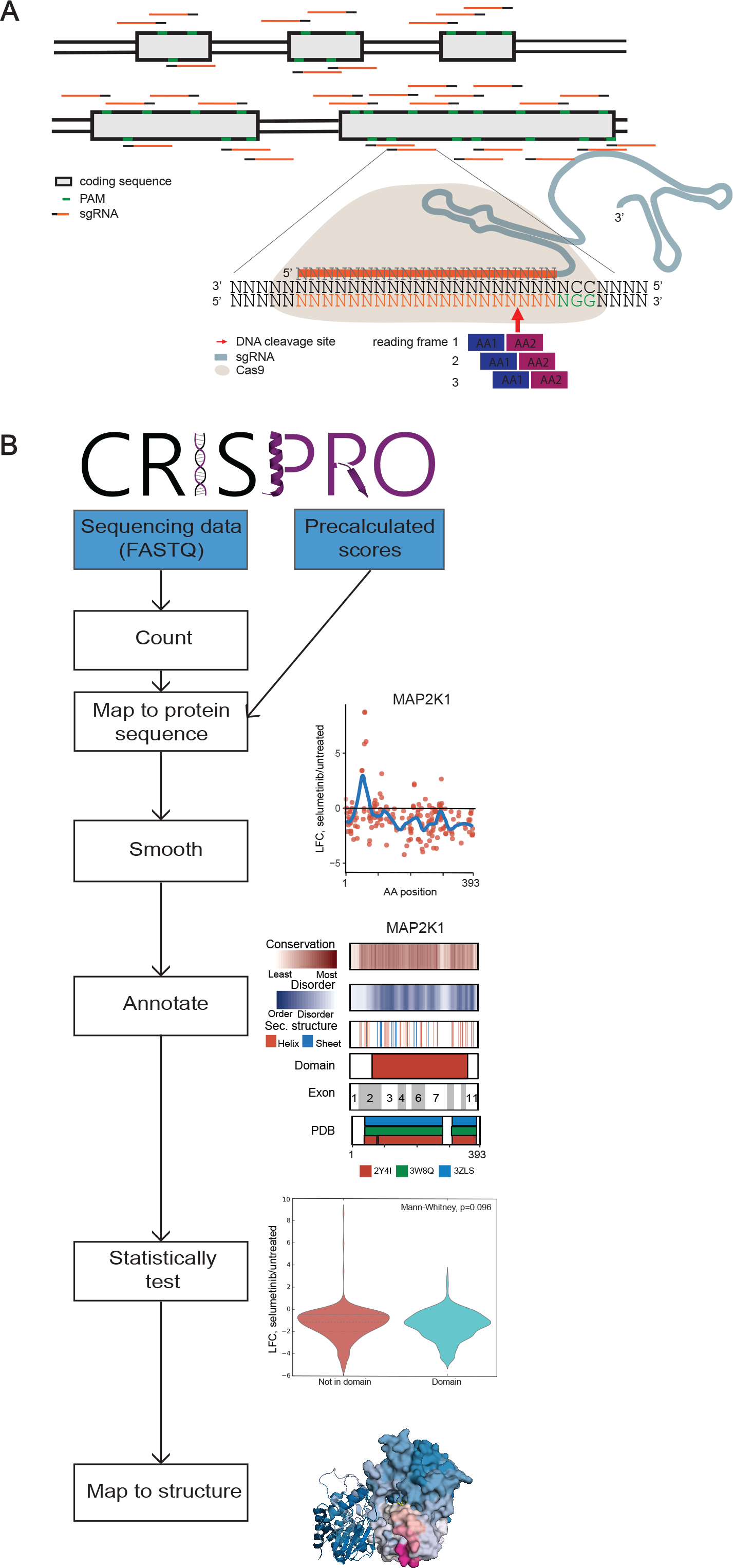
**A)** Dense mutagenesis of protein coding sequence by pooled CRISPR screening approach. Single guide RNAs target every possible PAM in the coding sequence of a gene. Guide RNAs are mapped to the two amino acids closest to the Cas9 cleavage site. **B)** Overview of the CRISPRO pipeline. Two input options are either FASTQ files or a precalculated score file (blue). PDB ID structure: 4MNE.

To gain mechanistic insights into genetic dependencies, we leverage CRISPR tiling screens, protein and nucleotide sequence level annotations, and 3D visualization of protein structure to elucidate functional residues and predict phenotypic outcome of genome editing in a singular computational pipeline, CRISPRO. To test and develop CRISPRO we first re-analyzed previously published data by Munoz et al. [7]. This study described a set of dense mutagenesis CRISPR screens to identify cancer cell genetic dependencies, and to investigate the importance of guide RNA positioning in gene inactivation in 3 different cell lines. Subsequently we re-analyzed CRISPR tiling data from Donovan et al. [8] on *MAP2K1* and *BRAF* as a test. Additionally we validated the analytic and predictive power of CRISPRO with prospective dense mutagenesis CRISPR data for *ZBTB7A* and *MYB* [5,9]. We observe that amino acid sequence conservation, predicted intrinsic protein disorder, and domain structure are highly predictive of the functional requirement of protein sequences. These analyses nominate discrete protein sequences as essential for specific biological phenotypes. Furthermore, we demonstrate the flexibility of the CRISPRO pipeline analyzing orthogonal dense mutagenesis datasets such as ectopic saturation mutagenesis. Finally, we derived a machine learning based model based on CRISPRO features to predict guide RNA efficacy in loss-of-function screens, providing improved predictive performance compared to tools primarily utilizing nucleotide features. The CRISPRO tool is freely available as open source software along with sample datasets at http://gitlab.com/bauerlab/crispro.

## Results

### Development of the CRISPRO tool

CRISPRO inputs next generation sequencing datasets resulting from dense mutagenesis CRISPR screens and maps functional scores associated with guide RNAs to genome, transcript, and protein coordinates. Briefly, each guide RNA is mapped to the two codons adjacent to the Cas9 cleavage site (see Methods) (Figure 1A). Smoothing via LOESS regression is performed in order to model local trends of the CRISPR perturbation effect over the entire protein and to provide scores for amino acids with no assigned guides. Calculating individual scores for guide RNAs is coupled with visualization of functional scores and tracks containing domain structure (InterPro [10]), secondary structure prediction, disordered region prediction and PROVEAN functional predictions based on interspecies conservation [11–18]. Visualization is also available at the tertiary structure level, where peptide fragments are aligned to existing protein structures in the Protein Data Bank (PDB, www.rcsb.org) and recolored in a heatmap style reflecting functional scores of amino acid residues [19] (Figure 1B). These functionally annotated structures may identify critical interfaces between the analyzed protein and other biomolecules as well as inform biophysical and chemical biology hypotheses.

When multiple genes are targeted in a CRISPR screen, CRISPRO defines hit genes with strong functional effect and performs analyses of correlation between functional scores and annotations. This is done pooled for the hit genes for each score as well as individually for each hit gene. To test the CRISPRO tool, we evaluated its performance with published datasets. Munoz et al. performed CRISPR pooled screening dense mutagenesis of 139 genes in 3 cancer cell lines. They reported guide RNA sequences with associated log_2_ fold change transformed by z-score for cellular dropout. This data was used as input for CRISPRO. Using default settings, CRISPRO defined 69, 52 and 77 hit genes for the DLD1, NCI-H1299, and RKO cell lines respectively (at least 75% of guides for a gene having a z-score less than 0, see Methods), largely overlapping the hit genes identified by Munoz et al. (Figure S1, Table S1). The user can optionally override the CRISPRO default hit gene calling and assign custom hit genes for analysis.

CRISPRO can also be used for calculation of functional scores per guide RNA (defined as log_2_ fold change between control and test condition) by using next generation sequencing (NGS) data as input. The tool includes an option to normalize guide RNA counts to a set of assigned negative control guide RNAs. When using NGS data as input, the tool outputs quality control metrics regarding the deep sequencing data.

### Association of genome editing functional outcome with conservation and disorder

Targeting amino acids in predicted protein domains has previously been shown to be associated with heightened CRISPR functional scores [4,7]. We confirmed with the Munoz et al. dataset that CRISPRO identified that guide RNAs targeting sequences inside as compared to outside domains showed more negative dropout scores (Figure 2A, S2A, D). In addition, several groups have previously shown that evolutionary conservation is correlated with CRISPR screen functional score [7,20]. We compared the CRISPR functional scores with the PROVEAN conservation scores. As expected, using the CRISPRO tool, we observed a correlation between conservation and functional scores across all 3 cell lines tested by Munoz et al (Spearman correlation, DLD1: *ρ*=0.24, p<0.001; NCI-H1299: *ρ*=0.3, p<0.001; RKO: *ρ*=0.29, p<0.001) (Figure 2B, S2B, E). These results are consistent with the hypothesis that targeting conserved as compared to nonconserved protein coding sequences is more likely to be associated with in-frame loss-of-function alleles. Comparing all the hit genes in the dataset, we observed a trend that correlation scores between conservation and CRISPR score were higher for genes at which the PROVEAN score had a larger standard deviation. This suggests that PROVEAN scores may be most predictive when they are widely distributed for a gene. In addition, genes that are more conserved (lower median PROVEAN score) tended to have lower median CRISPR score compared to less conserved genes, suggesting that PROVEAN score is not only predictive of the CRISPR score within a gene but also between different genes (Figure 2D, S2G,I).

**Figure 2.**
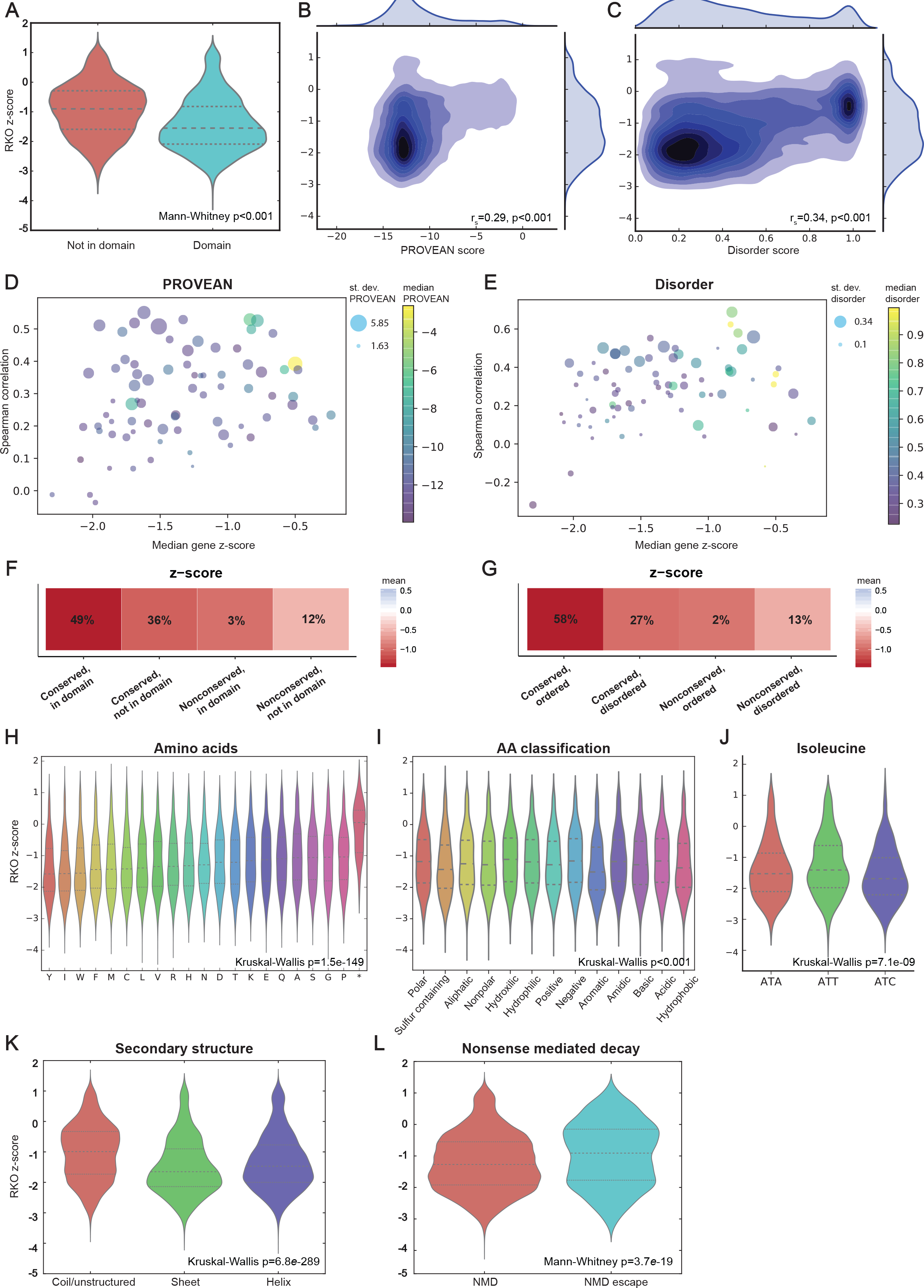
Correlation of functional scores for representative cell line RKO. **A)** Violin plot showing the distribution difference for guide RNA RKO z-scores targeting domains versus targeting outside of predicted domains (as defined by InterPro). **B)** Density plot showing the relation between RKO z-score and PROVEAN score (more negative is more conserved). **C)** Density plot showing the relation between RKO z-score and disorder scores (1 equals disorder, 0 equals order). **D)** Scatter plot showing the relation of median RKO z-score (x-axis), standard deviation (distribution) of PROVEAN score (marker size), and the median of the PROVEAN score (marker color) with the amount of correlation between PROVEAN scores and RKO z-scores (y-axis), for every gene. **E)** Analogous to D, but for disorder score in place of PROVEAN score. **F)** Heatmap showing the mean RKO z-score and the percentage guide RNAs falling into groups categorized based on domain annotation and conservation. **G)** Heatmap showing the mean RKO z-score and the percentage guide RNAs falling into groups categorized based on conservation and disorder score. **H)** RKO z-score distribution per amino acid. **I)** RKO z-score distribution per (non-mutually exclusive) amino acid class: polar (S, T, Y, N, Q), nonpolar (G, A, V, C, P, L, I, M, W, F), hydrophobic (A, V, I, L, M, F, Y, W), hydrophilic (S, T, H, N, Q, E, D, K, R), positively charged (R, H, K), negatively charged (D, E), aliphatic (A, G, I, L, P, V), aromatic (F, W, Y), acidic (D, E), basic (R, H, K), hydroxilic (S, T), sulfur containing (C, M), and amidic (N, Q). J) RKO z-score distribution per codon encoding for isoleucine (I). **K)** Distribution of RKO z-scores for guides targeting amino acids with different predicted secondary structure: coil/unstructured, sheet, or helix. **L)** Distribution for RKO z-scores for guides targeting sequences that are predicted to undergo or escape nonsense mediated decay (NMD).

We compared the effects of targeting domain annotated sequences to conserved sequences. We grouped guide RNAs based on both conservation (using PROVEAN score threshold -6) and domain assignment, resulting in four groups: 1) conserved, in domain; 2) conserved, not in domain; 3) nonconserved, in domain; 4) nonconserved, not in domain. Comparing the mean of these groups showed that amino acids in a domain and with a high conservation were associated with the greatest effect (most negative fitness scores). Notably, within the ‘not in domain’ groups, conserved residues had a more negative mean fitness score than those of nonconserved residues (Figure 2F, S2K, M).

In addition to conservation and domain assignment, we found that protein disorder score, a prediction of intrinsically disordered regions in proteins, also showed a correlation to functional CRISPR scores. The overall trend was that targeting a region with higher order (disorder score closer to 0) was associated with more profound functional score (Spearman correlation, DLD1: *ρ*=0.31, p<0.001; NCI-H1299: *ρ*=0.27, p<0.001; RKO: *ρ*=0.34, p<0.001) (Figure 2C, S2C, F). Similar to the finding for PROVEAN conservation scores, genes with wider distribution of disorder scores (higher standard deviation) demonstrated higher correlation with CRISPR scores compared to those with more narrowly distributed disorder scores. Also between genes, those with higher predicted order had higher negative median dropout scores as compared to those with higher predicted disorder (Figure 2E, S2H, J). We tested the relationship between disorder and conservation by grouping guides scores in 4 categories: 1) conserved, ordered; 2) conserved, disordered; 3) nonconserved, ordered; 4) nonconserved, disordered (Figure 2G, S2L, N). We found the most negative fitness scores for guides targeting conserved and ordered positions. This suggests that conservation and disorder can be used to further refine the set of key functional residues within a protein. Notably, we observed very few guides in the nonconserved and ordered category.

Association of genome editing functional outcome with protein primary and secondary structure We evaluated the impact of amino acid identity at the cleavage site as compared to guide RNA dropout score across the three cell lines. Amino acids with largest effect scores across the cell lines were tyrosine (Y), tryptophan (W), methionine (M), isoleucine (I), and leucine (L) (median scores for these in DLD1 < −1.25, Kruskal-Wallis: p=3e−136; NCI-H1299 < −1.7, Kruskal-Wallis: p=1.1e-93; RKO < −1.39, Kruskal-Wallis: p=1.5e-149) (Figure 2H, S3H, J). Selenocysteine (U) also showed a strong effect, however, this rare amino acid was only found twice in the screen and therefore was excluded from further analysis. Tyrosine and tryptophan are the heaviest amino acids (˜181 and 204 Da), and we hypothesized that their deletion might especially impact protein folding. They are also hydrophobic, as are methionine and isoleucine, which play a driving role in folding of the protein core [21]. Amino acids were then classified into 13 non-mutually exclusive groups: polar (S, T, Y, N, Q), nonpolar (G, A, V, C, P, L, I, M, W, F), hydrophobic (A, V, I, L, M, F, Y, W), hydrophilic (S, T, H, N, Q, E, D, K, R), positively charged (R, H, K), negatively charged (D, E), aliphatic (A, G, I, L, P, V), aromatic (F, W, Y), acidic (D, E), basic (R, H, K), hydroxilic (S, T), sulfur containing (C, M), and amidic (N, Q). This classification demonstrated more negative CRISPR scores for guide RNAs targeting hydrophobic amino acids as well as the partially overlapping groups of aromatic and sulfur containing amino acids (Figure 2I, S3I, K, S4). We tested if the reason for more negative scores at methionine might be due to targeting the start codon, but methionine at the start position of a protein sequence did not show a significantly different fitness score than methionine throughout the rest of the protein in any of the tested cell lines (Mann-Whitney-U test, DLD-1: p=0.229; NCI-H1299: p=0.161; RKO: p=0.431) (Figure S5).

We tested if the impact of disrupting individual codons could be due to the nucleotide identity of the codon itself rather than the encoded amino acid. If the functional effect were solely dependent on the amino acid, different codons for the same amino acid should have a similar score distribution. The only difference in average z-score comparing different codons for the same amino acid was observed for isoleucine (Kruskal-Wallis, DLD1: p=6e-13; NCI-H1299: p= 9.5e-05; RKO: p<0.001) (Figure 2J, S3L, M), where codon ATC had more negative dropout scores than codons ATT and ATA in all three cell lines. Previous data have suggested ATC may have enhanced translation as compared to other codons of isoleucine and may therefore influence protein folding [22,23]. All other amino acids had similar score distributions for all codons.

In addition to these amino acid classifications we predicted a consensus secondary structure by amalgamating the results of several publicly available tools (see Methods for details). We found that guide RNAs targeting sequences predicted to have helix or sheet secondary structure have a greater effect than those targeting sequences predicted to have a coil secondary structure or no secondary structure (Figure 2K, S3B, E).

### Association of genome editing functional outcome with mRNA annotations

Nonsense mediated decay (NMD) is the expected result of the introduction of a premature termination codon (PTC) by a frameshift indel following CRISPR/Cas9 cleavage repair. Exon-junction complex (EJC)-mediated NMD follows the 50 nucleotide rule, meaning that if a PTC resides more than 55 nucleotides upstream of the last exon-exon junction, the terminating ribosome will fail to remove the EJC, causing EJC-mediated NMD. Thus, guide RNAs targeting less than 55 nucleotides upstream of the final exon-exon junction should be able to escape NMD, whereas all others are sensitive to this quality control process [24]. We find that when applying this rule, guide RNAs targeting sequences with the ability to escape NMD indeed have less effect on the functional score (Mann-Whitney-U, DLD1: p=2.2e-37; NCI-H1299: p=1.8e-08; RKO: p=3.7e-19) (Figure 2L, S3C, F). These results suggest that triggering NMD is a major mechanism of genome editing induced loss-of-function alleles (Figure S11B).

We evaluated predictive value of some other mRNA-level annotations, including propensity for exon skipping, distance to exon-intron junction, and fraction of transcript isoforms targeted. We hypothesized that exons that were multiples of 3 would be less functionally essential as compared to those that were not multiples of 3, since mutations could produce mRNA with intact reading frame [25]. It has been reported that exon skipping can be caused (besides alternative splicing) by both point mutations and by CRISPR-induced indels [26]. We were not able to observe a pervasive impact of exon skipping on CRISPR score, with no significant difference in dropout phenotypes between guide RNAs targeting multiple-of-3 as compared to other exons (Figure S3A, D, G). We hypothesized that cleavage sites adjacent to exon-intron borders might have heightened functional scores since they could perturb splice sites in addition to protein coding sequences. However, we were unable to detect a significant difference in guide RNA dropout score for guides targeting close to as compared to distant from exon-intron borders (Figure S6A, B, D, E, G, H). We hypothesized that targeting sequences shared among transcript isoforms would be more effective than targeting unique isoforms. We observed that the fraction of targeted transcripts only makes a modest difference in CRISPR scores (Spearman correlation, DLD1: *ρ*=0.068, p<0.001; NCI-H1299: *ρ*=0.054, p<0.001; RKO: *ρ*=0.084, p<0.001) (Figure S6C, F, I).

### Association of genome editing functional outcome with nucleotide annotations

Several tools exist to predict either the on-target activity of guide RNAs, which can be defined as the likelihood of creating an indel at a given locus, such as the Doench (2016, Rule Set 2) score, Moreno-Mateos score, and the Wong score, among others [27]. In case of CRISPR experiments utilizing a U6 promoter to express the guide RNA, the Doench score has been shown to have the best performance among the publicly available on-target predictors [27]. Therefore, we focused on the Doench score in our analyses. The Doench score utilizes nucleotide and spacer features like melting temperature without explicitly including protein level features [28]. For CRISPR scores from the Munoz et al. dataset, we found that the Doench score was correlated with observed CRISPR score (Spearman correlation, DLD1: *ρ*=0.26, p<0.001; NCI-H1299: *ρ*=0.25, p<0.001; RKO: *ρ*=0.18, p<0.001) (Figure S7A, D, G) [28].

Additionally, we tested predicted frameshift scores with guide RNA score. We hypothesized that guide RNAs more likely to produce frameshift as compared to in-frame alleles would be associated with a greater effect on phenotypic score. However, we did not detect any association between the out-of-frame score [29] (as retrieved from the CRISPOR database) with the phenotypic CRISPR scores (Figure S7B, E, H)

### Linear maps of genome editing functional outcomes

CRISPRO outputs linear tracks showing functional CRISPR scores on a per guide RNA basis. Also a LOESS regression is performed between guide RNA functional scores and protein primary sequence location. LOESS regression parameters were calibrated by the length of the protein and the assumption that guide RNAs were uniformly distributed throughout a protein. This allows for plotting of interpolated scores for amino acids that are not targeted by a guide RNA. Several protein-level functional annotations are plotted below the guide RNA scores. These include PROVEAN conservation scores, disorder scores, secondary structure predictions, InterPro domain annotations [10], and aligned structures available from the PDB. These linear maps are generated for every gene included in the analysis, providing a visual overview of the data and enabling identification of potential regions of interest within a protein at a glance. For example, for *PLK1* and *AURKA* (Figures 3A, 3B) the largest negative impact of guide RNAs on cellular fitness is observed at conserved, ordered positions, with secondary structure predictions, and at domains. Reciprocally the least negative impact on cellular fitness is found at regions with high disorder, little conservation, lack of secondary structure, and without domain annotation. *CTNNB1* (Figure 3C) is a strong hit gene in only one of the three cell lines tested by Munoz et al., DLD1. However, in this cell line there is agreement between the most negative phenotypic CRISPR scores and conservation, disorder, secondary structure, and domain annotation.

**Figure 3.**
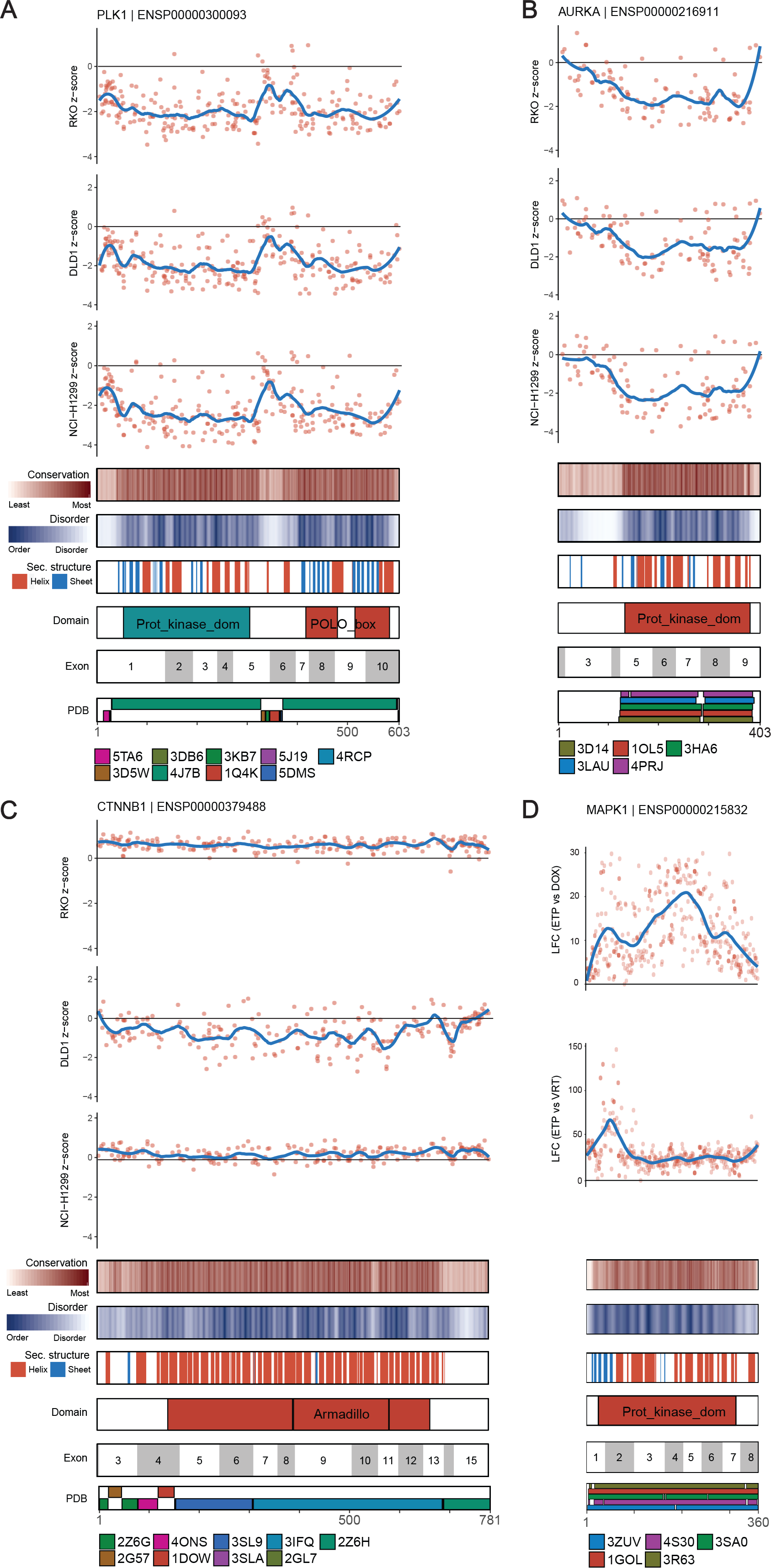
Linear maps. **A)** PLK1, cell fitness scores from the CRISPR dataset of Munoz et al.. **B)** AURKA, cell fitness scores from the CRISPR dataset of Munoz et al. **C)** CTNNB, cell fitness scores from the CRISPR dataset of Munoz et al.. **D)** *MAPK1*, log_2_ fold change (LFC) of MAPK1/ERK2 mutant abundance following DOX induction, relative to mutant abundance in the early time point (LFC (ETP vs. DOX)) and LFC of *MAPK1/ERK2* mutant abundance following DOX induction in the presence of 3 μM VRT-11E relative to mutant abundance in the early time point (LFC (ETP vs. VRT)), from the dataset of Brenan et al..

The linear mapping functionality of CRISPRO can be readily extended to non-CRISPR datasets. We used CRISPRO to visualize data produced by ectopic saturation mutagenesis of *MAPK1/ERK2* as performed by Brenan et al. [30]. This study tested the function of almost all possible *MAPK1/ERK2* missense mutations to identify gain-of-function and loss-of-function alleles. In this A375 cell line system loss-of-function MAPK1 mutants are associated with more rapid proliferation [30]. We summed functional scores for every amino acid substitution at a given position and normalized the summed scores to have a minimal positional score of 0. This resulted in two normalized datasets, with the log_2_ fold change (LFC) of *MAPK1* mutant abundance following DOX induction relative to an early time point (ETP vs DOX) to find loss-of-function alleles or in presence of VRT-11E, a small molecule ERK1/2 kinase inhibitor (ETP vs VRT) to find drug-resistance alleles (Figure 3D). The linear map generated by CRISPRO shows loss-of-function mutants at various sequences with high conservation and low disorder, whereas the drug resistance alleles are concentrated at the ATP-binding pocket around residues 25 to 70 [30] (Figure 3D). These data illustrate how CRISPRO can be used to flexibly map a variety of functional scores to protein annotations.

### Visualizing genome editing functional outcomes with protein structures

To further develop structure-function hypotheses from dense mutagenesis data, CRISPRO maps calculated functional scores to protein structures. CRISPRO uses BLAST [31] to search the PDB for all available protein structures and optionally downloads additional structures defined by the user. CRISPRO aligns the structures to the protein sequence and uses PyMOL [32] to recolor the structure based on CRISPR scores (see Methods). By default, CRISPRO sets a two-color heatmap based on the distribution of scores in the dataset such that the more extreme of the 5%ile or 95%ile guide RNA score demarks the last bin and the heatmap is centered around 0 (Figure S8). Per above, for PLK1 the lowest fitness scores were observed in the protein kinase and polo box domains. For both these domains, protein structures are available in the Protein Data Bank (PDB IDs 5TA6, 3FVH), onto which the interpolated CRISPR scores were mapped. The protein kinase domain structure 5TA6 shows the competitive inhibitor 5,6-dihydroimidazolo[1,5-f]pteridine binding at the ATP-binding pocket [33]. The noncatalytic polo box domain structure 3FVH shows the phosphothreonine mimetic peptide Ac-LHSpTA-NH2 binding at a key protein-protein interaction site [34]. Extremely low fitness scores were observed adjacent to these ligand binding sites, demonstrating the capacity of CRISPRO 3D mapping to highlight important protein regions (Figure 4A, B).

**Figure 4.**
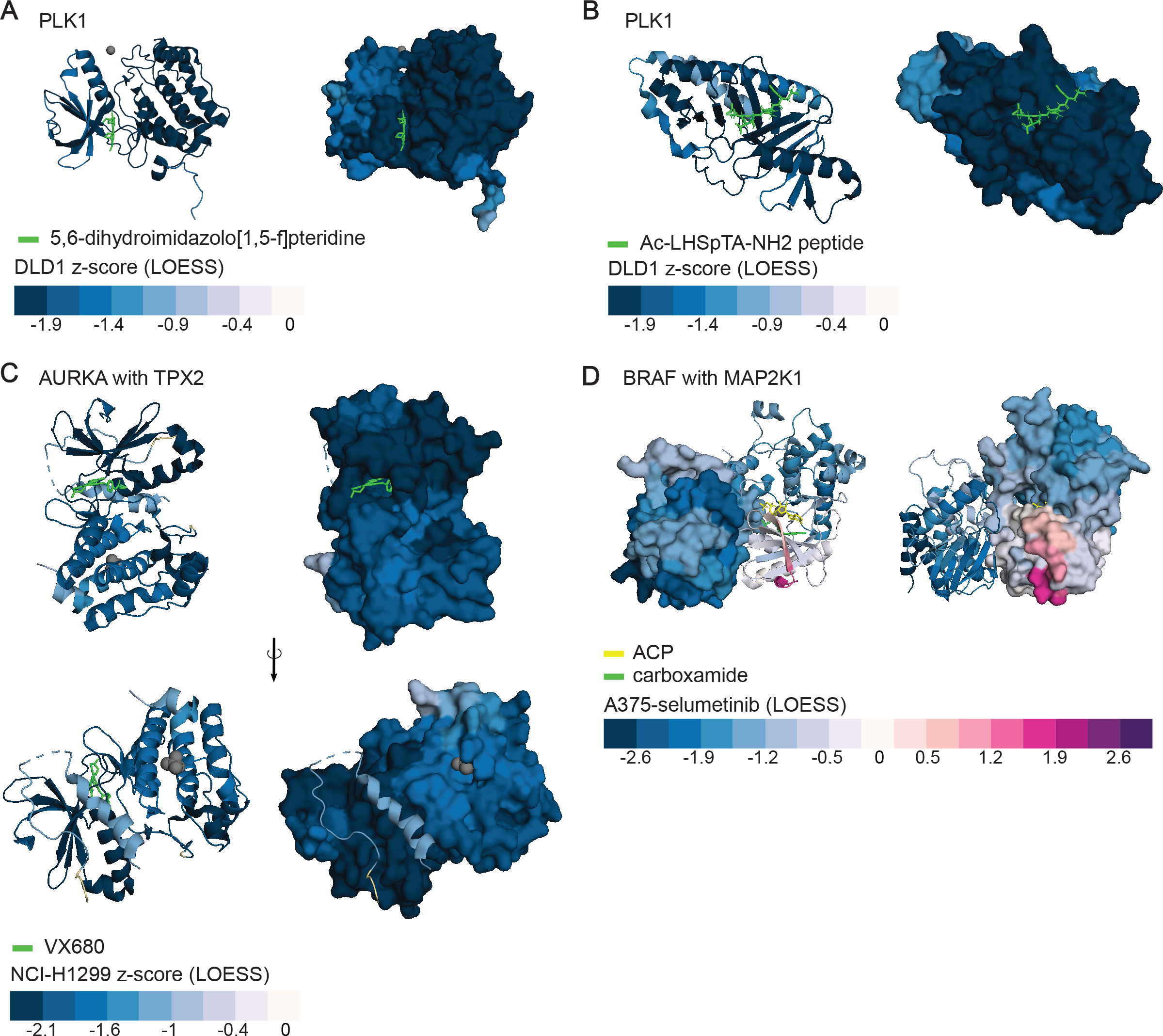
3D structure maps. **A)** PLK1, PDB ID: 5TA6. Mapped scores are DLD1 z-score (LOESS interpolation) of PLK1 (protein kinase domain, AA37-330, cartoon presentation in the left panel, surface presentation in the right panel) in complex with 5,6-dihydroimidazolo[1,5-f]pteridine inhibitor (green). Zinc ion is displayed as a grey sphere. **B)** PLK1, PDB ID: 3FVH. Mapped scores are DLD1 z-score (LOESS interpolation) of PLK1 (polo box domain, AA368-604) in complex with Ac-LHSpTA-NH2 peptide. Both surface (right) and cartoon (left) presentation. **C)** AURKA with TPX2. PDB ID: 3E5A. Mapped scores are NCI-H1299 z-score (LOESS interpolation) of AURKA (presented as surface in left panels, right as a cartoon, AA125-389, protein kinase domain) and TPX2 (presented solely as cartoon, AA6-21, 26-42, Aurora-A binding domain) in complex with VX680, an ATP-competitive small molecule inhibitor. Sulfate ion is displayed as a grey sphere. **D)** BRAF and MAP2K1. PDB ID: 4MNE. Mapped scores A375 selumetinib (LOESS interpolation) of BRAF (surface in left panel, cartoon in right, AA449-464, 469-722, protein kinase domain) and MAP2K1 (cartoon in left panel, surface in right, AA62-274, 307-382, protein kinase domain). Ligands ACP in yellow, and 7-fluoro-3-[(2-fluoro-4-iodophenyl)amino]-N-{[(2S)-2-hydroxypropyl]oxy}furo[3,2-c]pyridine-2-carboxamide in green. Magnesium ion is displayed as a grey sphere.

Another example of CRISPRO to highlight regions of 3D interaction with both small molecule inhibitors of catalytic active sites as well as with sites important for protein-protein interactions is found at AURKA, which is a member of a family of kinases that control progression through mitotic cell division [35]. The 3E5A structure shows AURKA in complex with both TPX2, a protein that serves as an allosteric activator of AURKA, as well as with VX680, an ATP-competitive small molecule inhibitor of kinase activity. Both the interaction sites of AURKA with TPX2 and AURKA with VX680 showed extremely low fitness scores (Figure 4C, S9). These results confirm that CRISPRO’s analyses and visualization can indicate functional regions of a protein and suggest it could help to identify potential regions of interest for chemical biology.

We used CRISPRO to map to available protein structures the results of the CRISPR screen of *MAP2K1* and BRAF in presence of a MEK inhibitor selumetinib to identify drug-resistance alleles [8] (Figure S10A, B). Although no structures of MAP2K1 with selumetinib were available, the structure PDB ID 4MNE shows the allosteric inhibitors ACP and carboxamide which are thought to occupy the same binding pocket as selumetinib (Figure 4D). The positive CRISPR phenotypic scores, indicating position of drug-resistance alleles (mapped in purple), showed that these positions are adjacent to the site of small molecule inhibitor binding, whereas other regions of MAP2K1 distant from small molecule binding only showed negative phenotypic scores, consistent with negative fitness effect from MAP2K1 loss-of-function. Moreover, BRAF, which does not directly bind to the small molecule inhibitors, only showed negative fitness scores, with some of the most negative scores concentrated at the BRAF:MAP2K1 protein-protein interaction interface. Overall these results demonstrate the capacity of the mapping function of CRISPRO to identify critical protein interfaces for functional small molecule active site or allosteric interactions, or sites of protein-protein interactions.

### Prediction of genome editing functional outcome

Given that various CRISPRO features such as conservation, disorder, and Doench score were correlated with CRISPR scores, we sought to test if the collection of features and annotations used in CRISPRO could be used to predict guide RNA efficacy in phenotypic screens. Before fitting a model, we preprocessed the feature set and target variable. The target variable was the average CRISPR score for a guide RNA in the three cell lines from the Munoz dataset. Then, we filtered out genes with a mean average CRISPR score > −1 to remove potential noise from genes with marginal phenotype. Next, we scaled the average CRISPR score by gene to make features and target variables comparable across genes. Regarding the feature set, we used protein, mRNA, and guide annotations, along with nucleotide positional features (see Methods). In total, the training data consisted of 10,286 guide RNAs from 42 genes. Using a simple lasso model we observed an encouraging predictive power (R^2^ = 0.35, Spearman *ρ* =0.59 in 10 Group Fold Cross Validation) and that disorder and PROVEAN scores were the most predictive features followed by the Doench score.

Next, to elucidate potential nonlinear interactions among features and to build a more robust model, we applied gradient boosting decision trees to the CRISPRO annotation set with nucleotide features incorporated. Parameters for the gradient boosting model were determined via 10 Group Fold Cross validation and the fitted model had greater R^2^ of 0.39 and Spearman *ρ* of 0.62 in 10 Group Fold Cross Validation and improved predictive performance on all test genes relative to Doench score alone (Figure 5A). Similar to the linear model, the most important features were the PROVEAN score of the targeted amino acid, Doench score of the guide RNA, disorder score of the targeted amino acid. Interestingly, this model recovered the identity of the last dinucleotide of the guide RNA as the fourth most important feature (Figure S11A). In order to observe the individual effects of various features in the model, partial dependency plots were used. The PROVEAN score, Doench score, and disorder score exhibited monotonic relationships in relation to the predicted scaled guide RNA score, with decreased guide RNA potency with increasing PROVEAN score and disorder score, and increased guide RNA potency with an increased Doench score (Figure S11E-I). Moreover, for the last dinucleotide of the spacer, GC and CG were associated with less effective guide RNAs relative to other dinucleotide pairs in the model (Figure S11C, D).

**Figure 5.**
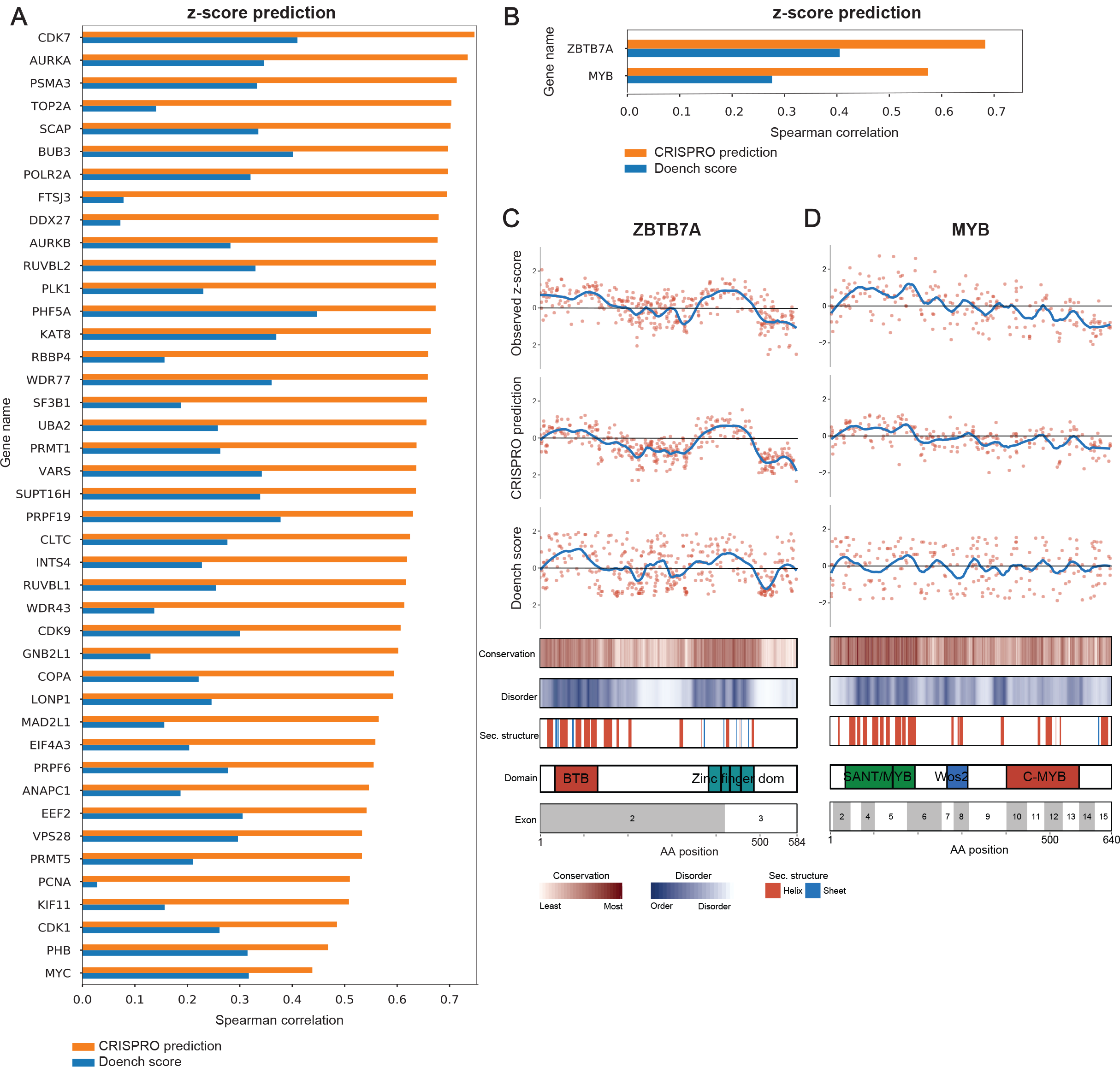
**A)** Spearman correlation between scaled average CRISPR z-score and predicted guide RNA efficacy by CRISPRO or Doench Rule Set 2 Score. CRISPRO prediction scores were generated by 10 group fold cross validation, where each gene was in a test set for prediction. **B)** Spearman correlation between scaled fitness scores and predicted guide RNA efficacy by CRISPRO or Doench Rule Set 2 Score at an independent dataset, prospectively generated and not used to train the model. **C-D)** Scatter plots of scaled observed guide RNA scores, CRISRPO prediction scores, and Doench scores, with LOESS regression shown by blue lines compared to position in protein. Protein level and mRNA level annotations are aligned underneath.

Importantly, we evaluated the gradient boosting model’s performance on an orthogonal dataset from a saturating mutagenesis experiment we performed evaluating the fitness impact of targeting two erythroid transcription factors, MYB and ZBTB7A, in erythroid precursor HUDEP-2 cells. The target variable, log fold change of guide RNA counts at the final time point, was transformed by z-score for each gene, analogous to the training data. The gradient boosting model increased the correlation of the predicted and actual values (R^2^ = 0.15, Spearman *ρ* =0.61) as compared to the correlation of the Doench score and actual values (Spearman *ρ* =0.34) (Figure 5B, C). These prediction results indicated by visual inspection critical functional regions at ZBTB7A and MYB, at the zinc finger cluster and SANT domain respectively, that were empirically validated (Figure 5C-D). These results indicate the utility of the prediction scores to forecast functional outcome of genome editing beyond the training dataset.

We have calculated CRISPRO prediction guide RNA scores across the hg19 proteome (available at gitlab.com/bauerlab/crispro). These predictions may help genome editing users select for functional studies guide RNAs likely to perturb the gene target.

## Discussion

The discovery of methods for programmable genome editing by CRISPR-Cas9 systems have offered unprecedented capabilities for comprehensive genetic perturbations in situ to investigate the sequence determinants of gene function. We have developed a widely-adaptable open-source computational tool, CRISPRO, to take deep sequence data from dense mutagenesis in situ pooled screens as input to compare functional scores with protein, transcript, and nucleotide level annotations, perform statistical association testing, and visualize functional results with linear maps and three-dimensional protein structures.

We confirmed prior observations that protein level annotations such as domain structure and interspecies sequence conservation help predict the functional outcome of CRISPR perturbation. Furthermore we demonstrate that other protein annotations such as disorder score have additional predictive utility.

By automatically mapping the phenotypic scores onto linear and 3D maps, the tool implicates discrete protein regions in specific biological phenotypes. Especially when combined with orthogonal genetic and biochemical data, the ensuing hypotheses may be prospectively tested to improve understanding of protein structure-function relationships and suggest critical interfaces as opportunities for rational targeting for bioengineering or therapeutics.

Beyond protein level annotations, we observed that transcript level (for example, NMD escape and isoleucine codon usage) and nucleotide level (for example, dinucleotide identity and Doench score) annotations offer additional layers of predictive power. We used these annotations to develop predictive models of genome editing functional outcomes by gradient boosting. We show improved performance as compared to previous models such as Doench score. We prospectively tested the predictions on an independently derived dataset, and validated the heightened predictive power of the CRISPRO informed predictions. We have generated prediction scores across all protein coding sequences.

The CRISPRO tool is flexible to incorporate additional annotations. We anticipate that inclusion of other annotations at various levels, including protein, transcript, chromatin, DNA sequence, and guide RNA, could further increase predictive power of the tool.

A current limitation of Cas9-mediated dense mutagenesis in situ is that the resolution is restricted by both the targeting range constraints of PAM sequence (such as NGG availability for SpCas9) and the variable and difficult to predict end-joining repair indel spectrum following nuclease cleavage dependent on nuclease, guide RNA, and target DNA, chromatin and cellular contexts. However with rapid advances in genome editing technology, the targeting range problem may be partially addressed by use of orthologous and engineered Cas nucleases with alternative PAM restriction, such as the recently described xCas9 with NGN PAM [36]. Ability to predict genome editing outcomes may improve with added knowledge of DNA repair determinants and empiric genome editing allele datasets. Furthermore non-nuclease genomic perturbation options continue to increase, such as the development of C and A base editors [37,38]. Since the CRISPRO tool is flexible with regard to input data, the resolution of its visualizations and predictive power of its associated annotations will likely only increase as genomic perturbation resolution continues to improve.

Although CRISPRO has been implemented as a tool to aid analysis and prediction of coding sequence perturbations, analogous inclusion of annotations from DNA and chromatin modifications, evolutionary conservation, genetic association studies, and other data types might ultimately be applied to the analysis and prediction of noncoding sequence perturbations as well.

## Conclusions

Here we describe CRISPRO open source software for the analysis of dense mutagenesis in situ pooled CRISPR screen datasets. We demonstrate the utility of various protein, transcript, and nucleotide level annotations to predict functional outcome of genome editing. The linear and 3D maps produced by CRISPRO may be used to develop hypotheses regarding structure-function relationships within mutagenized genes. CRISPRO annotations and models improve prediction of genome editing functional outcome.

## Methods

### CRISPRO pipeline

The CRISPRO pipeline is written completely in Python (The Python Software Foundation, https://www.python.org/) and R [39]. CRISPRO requires Python 2.7 and R>=3.4.1. Packages needed in R are tidyverse (ggplot2, dplyr, lazyeval, gridExtra, purr, RColorBrewer, readr), and DESeq2 (optional, when calculating scores). Package dependencies in Python are pandas (version >= 0.21.0), numpy, seaborn, matplotlib (version 1.5.3), PyMOL (version >= 2.1.0), scipy, and biopython.

There are two entry points to the CRISPRO pipeline. Users can either upload next generation sequencing data (sequence read files) in the FASTQ format or scores that have been calculated or precomputed (based on guide RNA or sequence coordinates in combination with the peptide ID).

The overview of the complete pipeline, from input to counting, mapping, annotating, testing and finally displaying the data onto structure, is displayed in Figure 1B. CRISPRO relies on a precompiled annotation set, which is available for publicly available for hg19. It will be possible for the user to compile other annotation datasets for different genome releases and organisms (e.g. hg38, mm10, etc.) with a script soon to be provided with the CRISPRO tool.

### Counting and mapping guides

The guide RNA counts for a sample are extracted from a given FASTQ file. To count the guides in each of the FASTQ files and to calculate functional scores, a list of identifiers, sample (condition) names and comparisons are needed. Identifiers can be either a list of genes, Ensembl peptide, transcript or gene IDs [40]. Guides are mapped to the protein sequence using information from the CRISPOR database [27]. This database contains all possible guides in the human genome (at coding exons), together with the genomic coordinate where they are predicted to cause a double strand break through Cas9 cleavage. Utilizing the CRISPOR database increases the speed of CRISPRO substantially since the mapping of guides can be precomputed. In addition, users do not have to provide guide sequences to count sequencing output.

To avoid the arbitrary decision of mapping a guide to one side of its cleavage site at both the nucleotide and amino acid levels, and given that CRISPR-Cas9 cleavage and NHEJ repair creates a typical indel spectrum affecting 1 up to 10 or more base pairs around the double strand break, each guide is mapped to the two amino acids nearest the double strand break by using genomic coordinates (Figure 1A).

Functional scores are calculated as the log_2_ fold change of the guide count in the sample groups provided and defined by the user. The user has the option to choose if the functional score is calculated by taking the average log_2_ fold change of replicates (ALFC method), or if the log_2_ fold change is calculated by using the DESeq2 R package [41]. DESeq2 is the preferred and default method of CRISPRO. It shrinks the value of the log_2_ fold change for a guide if read counts are low (noisy), to correct for the higher level of uncertainty. Reducing the fold change allows for confident comparison of all estimated fold changes across experiments.

### Off-target effect

Programmable nuclease mediated genomic cleavages can display modest negative fitness activity, presumably associated with activation of the DNA damage response. For this reason, we suggest it is best practice, especially in fitness/growth screens, that scores are normalized to functionally neutral genome targeting guides instead of non-targeting guides since non-targeting guides do not take into account nuclease-mediated gene-independent effects [42]. Guide RNAs targeting repetitive genomic sequences can have outsized non-specific negative fitness activity and may confound interpretation of perturbation screens [5]. Therefore, we suggest it is important to implement an off-target filter, to avoid high functional CRISPR scores solely caused by a high off-target effect (especially in fitness screens). We found in previous data (not shown) guides with a CRISPOR MIT off-target score lower than 5 often have extreme low fitness scores. Therefore, we include a default filter to remove any guide RNAs with CRISPOR MIT off-target score less than 5 [27,43]. This filter can be adjusted by the user.

### Smoothing

Scores for amino acids with no assigned guide RNA are interpolated via LOESS regression in the stats R package, using known guide scores and location as the model. LOESS regression is nonparametric, and uses weighted least squares to fit a quadratic curve on a contiguous subset of the data, in order to illustrate local trends of the CRISPR effect over the entire protein. The size of the subset of the data to which to fit a curve is determined by the span parameter, which is defined as 100 / protein length for a given protein. This parameter was set empirically based on correlation between the LOESS regression curve and other protein annotations such as PROVEAN and disorder scores (Figure S12). Moreover, the span parameter allows for approximately the same amount of data to be used to fit a local curve for various length genes with the assumption of uniform distribution of guide RNAs.

### Annotations

Essential to the analysis in the CRISPRO pipeline is annotating sequences and testing their correlation with calculated CRISPR scores. Sequences may influence CRISPR scores via effects at the DNA, RNA, or protein levels. At the DNA level, the target sequence and its surrounding context may specify guide RNA binding efficiency, off-target potential, or genomic repair preferences. Edits may affect mRNA splicing (by impacting cis-acting splice regulatory sequences), RNA stability (such as frameshifts that initiate nonsense mediated decay), or isoform usage (by targeting unique as compared to shared exons). At the protein level, the primary amino acid identity, secondary structures, likelihood of disorder, presence in identified domains, or interspecies/intraspecies constraint may influence the impact of mutations. CRISPRO utilizes one precompiled database with annotations from several genome-wide databases.

Annotations from publicly available databases include CRISPOR (guide efficiency score (Doench ‘16 [28]), out-of-frame score and off-target score), InterPro (domains), APPRIS (protein principal isoform), and Ensembl (exons, peptide and coding sequences) [10,27,44]. Additionally, the CRISPRO database contains precomputed conservation scores (PROVEAN[15]), exon length, DSB distance to 3’ and 5’ exon borders, the location in the protein (protein fraction), the predicted ability to escape nonsense mediated decay (NMD) (when the guide RNA targets upstream of −55 bp from the final exon-exon junction), the fraction of targeted protein isoforms per gene, disorder score, and secondary structure prediction.

PROVEAN (Protein Variation Effect Analyzer) is a protein sequence variant predictor that not only predicts the effect of single amino acid substitutions, like other commonly used tools such as PolyPhen and SIFT, but also predicts the effect of deletions. Since CRISPR-Cas9 cleavage creates a spectrum of indels, CRISPRO uses the effect score for single amino acid deletions generated by PROVEAN as a measure of conservation. CRISPRO’s original database is designed for hg19 proteins from Ensembl release 90. Therefore, we computed all PROVEAN scores for this database.

As described above, the DSB coordinate for each guide is obtained from the CRISPOR database. During mapping the guides to their corresponding amino acids in a protein also the distance to both exon borders is calculated, based on protein (genomic) coding coordinates from hg 19 Ensembl, release 90 (start and end points per exon).

Disorder scores for CRISPRO are pre-computed with VSL2b, a length-dependent predictor [17,18]. Disorder score is a prediction of intrinsically disordered regions (and proteins), which also have been called intrinsically unstructured, natively unfolded, natively disordered, or highly flexible regions. The classic model for protein activity description is “sequence leads to structure leads to function”, which holds for a majority of proteins. Therefore, disorder scores might be an important component in a guide’s final functional score. It is important to note that studies have showed that intrinsically disordered proteins and regions are not to be dismissed as regions necessarily lacking function. Intrinsically disorder regions are involved in binding other proteins, small molecules and nucleic acids. They have also been found to have a variety of biological functions, like signal transduction and cell or transcriptional regulation [17,45].

Multiple tools, PSSpred, PSIpred, SPINE X and RaptorX, were used to build a weighted consensus secondary structure prediction [11–14,16]. Each tool provides a probability score for a predicted secondary structure (either strand (B), helix (H) or coil (C)). For each amino acid, these scores are added up per secondary structure, and divided by the sum of all the options. This gives the weighted predictive score per secondary structure, whichever is the highest determines which secondary structure is predicted.

Two BLAST searches are used to align and annotate all available protein structures in the RCSB Protein Data Bank (PDB)[19,31]. The first search is done with complete protein sequences of the entire genome. These hits and alignments are directly available in CRISPRO’s standard annotation set. The second search is done per protein domain, as defined by the SMART database, to expand the range of available structures and to include partial structure hits which might have been missed in the first round of BLAST. For both BLAST searches the cut-off value for identity is 0.7 and e-value is 0.05. The results of the second BLAST search (domain only) are separated in an additional annotation file, which is only used when a user includes the option to map functional scores to structures in CRISPRO. In that case, any additional structures available for a protein are aligned with Biopython pairwise2 local alignment (using blosum62 matrix, gap open penalty: −10, gap extension penalty: −0.5) [46]. The option exists for the user to pass extra PDB ids (which might not have been found by the automated BLAST search) and the corresponding protein ID as input for CRISPRO. These structures would also be aligned with Biopython pairwise2 (same variables).

### General quality control and statistical testing

As part of its standard output CRISPRO provides summary statistics, quality information, guide density, functional scores and annotations based on raw FASTQ sequencing files. For each FASTQ file used as input, the following is calculated: total reads, mapped reads, percentage mapped reads, Gini score (a measure of inequality of the distribution), mean reads per guide, standard deviation reads per guide, minimum reads per guide, 10^th^ percentile reads per guide, median reads per guide, 90^th^ percentile reads per guide, and maximum reads per guide. All these values contribute to the quality control of the sequencing data and its mapping. In addition, raw read counts per guide are saved for each of the sequencing files (samples) and a Pearson correlation test is performed comparing all sequencing files.

Guide density and average guide distance is calculated for each gene individually. Guide density is calculated by dividing the total number of guides in a protein by the total number of amino acids. The distance between each of the guides is based on the first amino acid in the sequence it maps to, which is then averaged for all guides in a protein. Guides are filtered based on detection in the sequencing data. In other words, if according to CRISPOR there was a possible guide targeting the protein coding sequence, it is only taken into account if it was actually detected in the sequencing files and a functional score is calculated.

Functional scores are normalized to a negative control if available by subtracting the median negative control guide RNA score from each guide RNA score. It is optional for the user to assign negative and positive controls as input for CRISPRO. Negative controls can either be nontargeting guides or neutral gene-targeting guides. The latter negative control is encouraged when possible to control for the expected effect of gene-independent genome targeting events. Positive control guide RNAs could be targeting genes with known high effect, such as guides targeting ribosomal genes in the case of negative selection screens.

For the calculated (negative control normalized) functional scores of each gene the mean, standard deviation, first quartile, median, third quartile, and the interquartile range (IQR) are given. Additionally, the earth mover’s distance is calculated, indicating the cost of turning the distribution of scores of the protein into the distribution of the negative control distribution. Operationally CRISPRO defines a test gene as a hit for a given score (i.e. showing an overall phenotype of potential biological interest) in the CRISPR screen by checking if the IQR of the score contains 0, which corresponds to the median of the distribution of the negative controls. If not, the gene is labeled as hit. We have found that a Mann Whitney test between guides targeting a gene and nontargeting controls leads to the classification of most test genes as hits, since the number of guides is usually high, so even small effect sizes may be statistically significant. This tendency to identify many genes as significant hits may be exaggerated with use of nontargeting guides as negative control as compared to neutral genes [42]. It is possible for the user to define gene hits as an input for CRISPRO, in which case the default of using the IQR will be overwritten. For the purpose of further statistical testing, the direction of the hit is assigned, labeling the hit gene as either positive (median > 0) or negative (median ≤ 0).

Afterwards plots are generated to show correlation between every annotation CRISPRO provides and the functional scores. For the categorical annotations these are violin or box plots, for continuous data these are scatter plots. These plots are generated for each score for all hit genes pooled and for the individual hit genes. Appropriate statistical tests for each annotation are also performed (Spearman correlation, Mann-Whitney test, Kruskal-Wallis test with SciPy module in Python [47]).

### Mapping CRISPR scores to protein structures

When the user chooses to map the functional CRISPR scores to protein structures, all structures that were found by BLAST in the PDB (as described above) are downloaded. In case there are specific structures the user wants to map, regardless if these were found in the standard BLAST search, the user has the option to pass the PDB IDs and the corresponding protein ID as input for CRISPRO. These structures will also be included in all other standard output for CRISPRO, like the figures presenting annotations (linear tracks) and overview tables. Every PDB structure that is found (and complies with before mentioned conditions of the BLAST search) and added by the user will be mapped, even if there are multiple structures available for the same (sub)sequence of a protein.

The amino acid sequence of the structure is saved via the molecular visualization program PyMOL [32], and aligned with the full protein sequence to which the scores were mapped originally (described above). Based on these alignments, raw input text files for PyMOL are written, containing a list with the CRISPR functional score values corresponding to each amino acid present in the structure. It might occur that a structure has a different sequence than the original protein sequence, in which case there may be mismatches between amino acids, amino acids missing, and also extra amino acids present in the structure. If there are amino acids in the structure that are different but are aligned to an amino acid in the original protein, the corresponding score is mapped. If there are extra amino acids in the structure which cannot be aligned, no data will be mapped (shown in yellow).

Structures are then recolored in PyMOL, by loading the CRISPR functional scores in the B-factor field. Based on these values, a bin and corresponding color is assigned to each amino acid in the structure. The standard CRISPRO color legend exists of either 17 or 9 bins, from blue to dark purple, centered on 0. To be able to visually compare proteins and to distinguish important regions, bin size and boundaries are determined for each functional CRISPR score (separately for both raw and LOESS regressed scores), over all the proteins in the dataset. Set as the upper and lower border of the outermost bins are either the 5^th^ or 95^th^ percentile (and its inverse) of the score distribution, whichever is farther from 0. Every score lower or higher than this value will fall into those outer bins. The rest of the bins are evenly sized between these borders, resulting in a scale centered on 0 (Figure S8).

The recolored structures are saved as PyMOL session files (.pse), so the user can open them in the desktop version of PyMOL, and adjust the orientation or visuals of the structure before saving an image.

### Score prediction

Data Processing. Each set of guide RNA scores (average CRISPR scores) for a gene was multiplied by −1, if the mean score of the guide RNAs was less than 0, and z-score normalized. Multiplying by −1 allows us to make predictions for guide RNAs targeting genes that increase or decrease a phenotype, such that a predicted high CRISPR score is interpreted as having the greatest effect on phenotype for that gene, regardless of direction. Having each guide RNA normalized to other guide RNAs targeting the same gene allows for features to be comparable across all genes, whereas raw LFC or total z-score may not be comparable for guide RNAs of different genes due to inherent differences in gene function. In the Lasso Regression Model, features were scaled to have a value between 0 and 1 to allow for facile interpretation of model coefficients.

Predictive Models. For Lasso Regression, LassoCV was used from the scikit Learn package in python with default parameters to determine the optimal alpha parameter via the default cross validation method [48]. LGBMREgressor, from the LightGBM package in Python was used for the gradient boosting algorithms described above [49]. The hyperparameter space for the gradient boosted decision trees was explored using GridSearchCV from the scikit learn package in Python [48], yielding the following parameters differing from the default: (’bagging_freq’: 1, ‘colsample_bytree’: 0.75, ‘learning_rate’: 0.1, ‘max_depth’: 3, ‘min_child_samples’: 32, ‘n_estimators’: 512, ‘max_bin’: 63 Cross validation was performed by leaving out guides targeting 10% of proteins in the full training set (42 proteins).

Features. Targeted amino acid 1 and 2, domain occupancy status (InterPro), length of targeted exon multiple of 3, ability to escape nonsense mediated decay, single nucleotide and dinucleotide positional identities within guide RNA spacer (i.e. identity of nucleotide at position 17 in spacer) were all used as categorical features (categorical features are not one hot encoded in LightGBM implementation of gradient boosting decision trees). Numerical features included PROVEAN deletion score of the targeted amino acids 1 and 2, position in the gene, predicted disorder score of amino acids 1 and 2, GC content of the 20mer guide, length of the targeted Exon, and off-target score of the guide RNA. GC content of the 20mer guide was computed by adding the number of observed “G”s and “C”s in the 20mer and dividing the sum by the length of the guide (20 bp).

### Saturating mutagenesis CRISPR/Cas9 fitness screen in HUDEP-2

HUDEP-2 cells constitutively expressing lenti-Cas9 were transduced with a lentiviral guide RNA library containing puromycin resistance. 24 hours post transduction, cells underwent selection and erythroid based differentiation protocol. After 12 days of culture, the genomic DNA was captured allowing for next generation sequencing (NGS) of the guide RNA construct as previously described [5]. The fitness score was defined as the log_2_ fold change of counts in the final time point over the counts in the lentiviral plasmid sample.

**Figure S1 Hit genes. A)** Venn diagrams for each cell line, showing assigned number of hits per method (CRISPRO and Munoz et al.), and their overlap.Munoz et al. originally defined hits by setting a cut-off at an average z-score of −0.4, defining 74 (DLD1), 65 (NCI-H1299), and 78 (RKO) hits. Out of the hits set by Munoz et al., CRISPRO identified 65, 52, and 74 hits from DLD1, NCI-H1299, and RKO, missing a few hits which had a mean just below the −0.4 cut-off, with a relatively large standard deviation. Hits unique to CRISPRO had average z-scores close to −0.4, but had a smaller distribution of scores (Supplementary Table 1). CRISPRO assigned 4 and 3 additional hits for DLD1 and RKO cell lines. **B-D)** Violin plots for hits uniquely defined by CRISPRO or Munoz et al., with two example hits identified by both methods (PLK1 and AURKB). For each cell line, **(B)** DLD1, **(C)** NCI-H1299, **(D)** RKO.

**Figure S2 Correlation of functional scores for cell lines DLD1 and NCI-H1299. A)** Violin plot showing the distribution difference for guide RNA DLD1 z-scores targeting domains versus targeting outside of predicted domains (as defined by InterPro). **B)** Density plot showing the relation between DLD1 z-score and PROVEAN score (more negative is more conserved). **C)** Density plot showing the relation between DLD1 z-score and disorder scores (1 equals disorder, 0 equals order). **D)** Violin plot showing the distribution difference for guide RNA NCI-H1299 z-scores targeting domains versus targeting outside of predicted domains (as defined by InterPro). **E)** Density plot showing the relation between NCI-H1299 z-score and PROVEAN score (more negative is more conserved). **F)** Density plot showing the relation between NCI-H1299 z-score and disorder scores (1 equals disorder, 0 equals order). **G)** Scatter plot showing the relation of median DLD1 z-score (x-axis), standard deviation (distribution) of PROVEAN score (marker size), and the median of the PROVEAN score (marker color) with the amount of correlation between PROVEAN scores and DLD1 z-scores (y-axis), for every transcript. **H)** See G, now for disorder and DLD1 z-score. **I)** see G, now for NI-H1299 scores. **J)** See G, now for disorder and NCI-H1299 z-scores. K) Heatmap showing the mean DLD1 z-score and the percentage guide RNAs falling into groups categorized based on domain annotation and conservation. **L)** See K, now for NCI-H1299 z-scores. **M)** Heatmap showing the mean DLD1 z-score and the percentage guide RNAs falling into groups categorized based on conservation and disorder score. **N)** See M, now for NCI-H1299 z-scores.

**Figure S3 A)** Distribution of DLD1 z-scores for guides targeting in exons that are a multiple of 3 or not. **B)** Distribution of DLD1 z-scores for guides targeting amino acids with different predicted secondary structure: coil/unstructured, sheet, or helix. **C)** Distribution for RKO z-scores for guides targeting nucleotides that are able to escape NMD or that are subject to it. **D)** Distribution of NCI-H1299 z-scores for guides targeting in exons that are a multiple of 3 or not. **E)** Distribution of NCI-H1299 z-scores for guides targeting amino acids with different predicted secondary structure: coil/unstructured, sheet, or helix. **F)** Distribution for NCI-H1299 z-scores for guides targeting nucleotides that are able to escape NMD or that are subject to it. **G)** Distribution of RKO z-scores for guides targeting in exons that are a multiple of 3 or not. **H)** DLD1 z-score distribution per amino acid. **I)** DLD1 z-score distribution per (non-mutually exclusive) amino acid class: polar (S, T, Y, N, Q), nonpolar (G, A, V, C, P, L, I, M, W, **F)**, hydrophobic (A, V, I, L, M, F, Y, W), hydrophilic (S, T, H, N, Q, E, D, K, R), positively charged (R, H, K), negatively charged (D, E), aliphatic (A, G, I, L, P, V), aromatic (F, W, Y), acidic (D, E), basic (R, H, K), hydroxilic (S, T), sulfur containing (C, M), and amidic (N, Q). **J)** NCI-H1299 z-score distribution per amino acid. K) NCI-H1299 z-score distribution per amino acid class (see (I)). **L)** DLD1 z-score distribution per codon encoding for isoleucine (I). **M)** NCI-H1299 z-score distribution per codon encoding for isoleucine (I).

**Figure S4 A-C)** Percentage of amino acids in each amino acid class per transcript (see figure S3I). (A) DLD1 cell line, (B) NCI-H1299 cell line, (C) RKO cell line. **D-F)** Median z-score for each amino acid class per transcript (D) DLD1 cell line, (E) NCI-H1299 cell line, (F) RKO cell line.

**Figure S5** Distribution of z-scores for guide RNAs targeting the amino acid methionine at the start position of the protein, or any other place in the protein. A) DLD1 cell line, **B)** NCI-H1299 cell line, **C)** RKO cell line.

**Figure S6 A)** Density plot showing the relation between DLD1 z-score and the distance of the guide RNA predicted cleavage site to the 5’ exon border. **B)** Density plot showing the relation between DLD1 z-score and the distance of the guide RNA predicted cleavage site to the 3’ exon border. **C)** Density plot showing the relation between DLD1 z-score and the fraction of targeted transcripts of a gene by a guide RNA. **D)** Density plot showing the relation between NCI-H1299 z-score and the distance of the guide RNA predicted cleavage site to the 5’ exon border. **E)** Density plot showing the relation between NCI-H1299 z-score and the distance of the guide RNA predicted cleavage site to the 3’ exon border. **F)** Density plot showing the relation between NCI-H1299 z-score and the fraction of targeted transcripts of a gene by a guide RNA. **G)** Density plot showing the relation between RKO z-score and the distance of the guide RNA predicted cleavage site to the 5’ exon border. **H)** Density plot showing the relation between RKO z-score and the distance of the guide RNA predicted cleavage site to the 3’ exon border. **I)** Density plot showing the relation between RKO z-score and the fraction of targeted transcripts of a gene by a guide RNA.

**Figure S7 A)** Density plot showing the relation between DLD1 z-score and the Doench score. **B)** Density plot showing the relation between DLD1 z-score and the out-of-frame score. **C)** Density plot showing the relation between DLD1 z-score and the off-target score of a guide RNA. **D)** Density plot showing the relation between NCI-H1299 z-score and the Doench score. **E)** Density plot showing the relation between NCI-H1299 z-score and the out-of-frame score. **F)** Density plot showing the relation between NCI-H1299 z-score and the off-target score of a guide RNA. **G)** Density plot showing the relation between RKO z-score and the Doench score. **H)** Density plot showing the relation between RKO z-score and the out-of-frame score. **I)** Density plot showing the relation between RKO z-score and the off-target score of a guide RNA.

**Figure S8 Score distributions** with color legend, outer bin borders presented by red dotted line. **A)** DLD1 z-score. **B)** DLD1 z-score LOESS regression. **C)** NCI-H1299 z-score. **D)** NCI-H1299 z-score LOESS regression. **E)** RKO z-score. **F)** RKO z-score LOESS regression.

**Figure S9** Linear map of TPX2. From top to bottom: Graph showing RKO z-scores (each guide RNA with corresponding score, mapped to protein sequence (red dot), and LOESS interpolation (blue line)). DLD1 z-scores. NCI-H1299 z-scores. Predicted conservation scores (PROVEAN). Predicted disorder scores. Predicted secondary structure consensus. Domain annotations (InterPro). Exon annotations. Initial PDB alignments.

**Figure S10 A)** Linear map of BRAF. From top to bottom: Graph showing A375 with selumetinib scores (each guide RNA with corresponding score, mapped to protein sequence (red dot), and LOESS interpolation (blue line)). A375 with vemurafenib scores. MELJUSO with selumetinib scores. Predicted conservation scores (PROVEAN). Predicted disorder scores. Predicted secondary structure consensus. Domain annotations (InterPro). Exon annotations. Initial PDB alignments. **B)** Linear map of MAP2K1.

**Figure S11 A)** Feature importance in gradient boosting model by information gain when a feature is used to split the training data. Positional nucleotide features are 0-indexed (i.e. nucleotide 0 is in position 1 of the spacer sequence, dinucleotide 0 corresponds to positions 1 and 2 of spacer, where position 20 is PAM proximal). **B)** Pairwise Spearman correlation coefficient for all numerical and binary features in CRISPRO training set. **C)** Partial dependency plot of feature dinucleotide 9 in gradient boosting model. Blue fill shows standard deviation of relative CRISPR score impact for each dinucleotide. Higher relative CRISPR score impact can be interpreted as higher loss of function capability. **D)** Partial Dependency contour plot depicting relationship between Dinucleotides 9 and 8 in trained gradient boosting model. **E-I)** Partial dependency plots of PROVEAN score 2, disorder score 2, and Doench score along with pairwise interactions through partial dependency contour plots.

**Figure S12 A)** Correlation of LOESS regression CRISPR scores and corresponding positional disorder scores as a function of the span parameter in LOESS regression. Span parameter defined as amino acid window / 100. **B)** Correlation of LOESS regression CRISPR scores and corresponding positional PROVEAN scores a function of the span parameter in LOESS regression.

## List of abbreviations

Clustered regularly interspaced short palindromic repeats (CRISPR)
Non-homologous end joining (NHEJ)
Base pair (bp)
Insertion and deletion (indel)
Nonsense mediated decay (NMD)
Single guide RNA (sgRNA)
Protospacer adjacent motif (PAM)
Protein Data Bank (PDB)
Fetal hemoglobin (HbF)
Premature termination codon (PTC)
Exon-junction complex (EJC)
Next generation sequencing (NGS)
Log_2_ fold change (LFC)
Early time point (ETP)
Partial Dependency contour Plot (PDP)

## Declarations

### Ethics approval and consent to participate

Not applicable.

### Consent for publication

Not applicable.

### Availability of data and material

The datasets supporting the conclusions of this article are available from following published articles:

Munoz DM, Cassiani PJ, Li L, Billy E, Korn JM, Jones MD, et al. CRISPR screens provide a comprehensive assessment of cancer vulnerabilities but generate false-positive hits for highly amplified genomic regions. Cancer Discov. 2016;6:900-13. DOI: 10.1158/2159-8290.CD-16-0178

Donovan KF, Hegde M, Sullender M, Vaimberg EW, Johannessen CM, Root DE, et al. Creation of novel protein variants with CRISPR/Cas9-mediated mutagenesis: Turning a screening by-product into a discovery tool. PLoS One. 2017;12:1-13. DOI: 10.1371/journal.pone.0170445

Brenan L, Andreev A, Cohen O, Pantel S, Kamburov A, Cacchiarelli D, et al. Phenotypic Characterization of a Comprehensive Set of MAPK1/ERK2 Missense Mutants. Cell Rep 2016;17:1171-83. DOI: 10.1016/J.CELREP.2016.09.061

The generated analysis is available in this article and its supplementary information files.

The source code for CRISPRO is freely available at http://gitlab.com/bauerlab/crispro. CRISPRO is platform independent, relies on PyMOL 2.1.0 and is written in Python and R.

### Authors’ contributions

MAC, CMT, PGS, LP and DEB conceptualized the pipeline. VACS and MAC designed and developed the pipeline with help from CMT and QY. MAC, FS, MCC, TM, DEB designed CRISPR fitness screen, which was performed by FS. Supervision of the complete project by LP and DEB. Original manuscript was written by VACS, MAC and DEB, and modified by QY and LP. All authors approved and read the final manuscript.

## References

1. Shalem O, Sanjana NE, Zhang F. High-throughput functional genomics using CRISPR-Cas9. Nat Rev Genet. 2015;16:299–311.

2. Doudna JA, Charpentier E. The new frontier of genome engineering with CRISPR-Cas9. Science. 2014;346:1258096.

3. Zhou Y, Zhu S, Cai C, Yuan P, Li C, Huang Y, et al. High-throughput screening of a CRISPR/Cas9 library for functional genomics in human cells. Nature. 2014;509:487–91.

4. Shi J, Wang E, Milazzo JP, Wang Z, Kinney JB, Vakoc CR. Discovery of cancer drug targets by CRISPR-Cas9 screening of protein domains. Nat Biotechnol. 2015;33:661–7.

5. Canver MC, Lessard S, Pinello L, Wu Y, Ilboudo Y, Stern EN, et al. Variant-aware saturating mutagenesis using multiple Cas9 nucleases identifies regulatory elements at trait-associated loci. Nat Genet. 2017;49:625–34.

6. Canver MC, Smith EC, Sher F, Pinello L, Sanjana NE, Shalem O, et al. BCL11A enhancer dissection by Cas9-mediated in situ saturating mutagenesis. Nature. 2015;527:192–7.

7. Munoz DM, Cassiani PJ, Li L, Billy E, Korn JM, Jones MD, et al. CRISPR screens provide a comprehensive assessment of cancer vulnerabilities but generate false-positive hits for highly amplified genomic regions. Cancer Discov. 2016;6:900–13.

8. Donovan KF, Hegde M, Sullender M, Vaimberg EW, Johannessen CM, Root DE, et al. Creation of novel protein variants with CRISPR/Cas9-mediated mutagenesis: Turning a screening by-product into a discovery tool. PLoS One. 2017;12:1–13.

9. Masuda T, Wang X, Maeda M, Canver MC, Sher F, Funnell APW, et al. Transcription factors LRF and BCL11A independently repress expression of fetal hemoglobin. Science. 2016;351:285–9.

10. Finn RD, Attwood TK, Babbitt PC, Bateman A, Bork P, Bridge AJ, et al. InterPro in 2017—beyond protein family and domain annotations. Nucleic Acids Res. 2017;45:D190–9.

11. Yan R, Xu D, Yang J, Walker S, Zhang Y. A comparative assessment and analysis of 20 representative sequence alignment methods for protein structure prediction. Sci Rep. 2013;3:2619.

12. Jones DT. Protein secondary structure prediction based on position-specific scoring matrices. J Mol Biol. 1999;292:195–202.

13. Buchan DWA, Minneci F, Nugent TCO, Bryson K, Jones DT. Scalable web services for the PSIPRED Protein Analysis Workbench. Nucleic Acids Res. 2013;41:W349–57.

14. Faraggi E, Zhang T, Yang Y, Kurgan L, Zhou Y. SPINE X: improving protein secondary structure prediction by multistep learning coupled with prediction of solvent accessible surface area and backbone torsion angles. J Comput Chem. 2012;33:259–67.

15. Choi Y, Sims GE, Murphy S, Miller JR, Chan AP. Predicting the Functional Effect of Amino Acid Substitutions and Indels. PLoS One. 2012;7:e46688.

16. Wang Z, Zhao F, Peng J, Xu J. Protein 8-class secondary structure prediction using conditional neural fields. Proteomics. 2011;11:3786–92.

17. Oates ME, Romero P, Ishida T, Ghalwash M, Mizianty MJ, Xue B, et al. D2P2: database of disordered protein predictions. Nucleic Acids Res. 2012;41:D508–16.

18. Peng K, Radivojac P, Vucetic S, Dunker AK, Obradovic Z. Length-dependent prediction of protein intrinsic disorder. BMC Bioinformatics. 2006;7:208.

19. Berman HM, Westbrook J, Feng Z, Gilliland G, Bhat TN, Weissig H, et al. The Protein Data Bank. Nucleic Acids Res. 2000;28:235–42.

20. Erard N, Knott SR V, Hannon GJ. A CRISPR Resource for Individual, Combinatorial, or Multiplexed Gene Knockout. Mol Cell. 2017;67:348–354.e4.

21. Balchin D, Hayer-Hartl M, Hartl FU. In vivo aspects of protein folding and quality control. Science. 2016;353:aac4354.

22. Bartoszewski RA, Jablonsky M, Bartoszewska S, Stevenson L, Dai Q, Kappes J, et al. A synonymous single nucleotide polymorphism in DeltaF508 CFTR alters the secondary structure of the mRNA and the expression of the mutant protein. J Biol Chem. 2010;285:28741–8.

23. Lazrak A, Fu L, Bali V, Bartoszewski R, Rab A, Havasi V, et al. The silent codon change I507-ATC->ATT contributes to the severity of the ΔF508 CFTR channel dysfunction. FASEB J. 2013;27:4630–45.

24. Lindeboom RGH, Supek F, Lehner B. The rules and impact of nonsense-mediated mRNA decay in human cancers. Nat Genet. 2016;48:1112–8.

25. Lalonde S, Stone OA, Lessard S, Lavertu A, Desjardins J, Beaudoin M, et al. Frameshift indels introduced by genome editing can lead to in-frame exon skipping. PLoS One. 2017;12:e0178700.

26. Mou H, Smith JL, Peng L, Yin H, Moore J, Zhang X-O, et al. CRISPR/Cas9-mediated genome editing induces exon skipping by alternative splicing or exon deletion. Genome Biol. 2017;18:108.

27. Haeussler M, Schönig K, Eckert H, Eschstruth A, Mianné J, Renaud J-B, et al. Evaluation of off-target and on-target scoring algorithms and integration into the guide RNA selection tool CRISPOR. Genome Biol. 2016;17:148.

28. Doench JG, Fusi N, Sullender M, Hegde M, Vaimberg EW, Donovan KF, et al. Optimized sgRNA design to maximize activity and minimize off-target effects of CRISPR-Cas9. Nat Biotechnol. 2016;34:184–91.

29. Bae S, Kweon J, Kim HS, Kim J-S. Microhomology-based choice of Cas9 nuclease target sites. Nat Methods. 2014;11:705–6.

30. Brenan L, Andreev A, Cohen O, Pantel S, Kamburov A, Cacchiarelli D, et al. Phenotypic Characterization of a Comprehensive Set of MAPK1/ERK2 Missense Mutants. Cell Rep. 2016;17:1171–83.

31. Camacho C, Coulouris G, Avagyan V, Ma N, Papadopoulos J, Bealer K, et al. BLAST+: architecture and applications. BMC Bioinformatics. 2009;10:421.

32. The PyMOL Molecular Graphics System. Version 2.0 Schrödinger, LLC.

33. Kiryanov A, Natala S, Jones B, McBride C, Feher V, Lam B, et al. Structure-based design and SAR development of 5,6-dihydroimidazolo[1,5-f]pteridine derivatives as novel Polo-like kinase-1 inhibitors. Bioorg Med Chem Lett. 2017;27:1311–5.

34. Yun S-M, Moulaei T, Lim D, Bang JK, Park J-E, Shenoy SR, et al. Structural and functional analyses of minimal phosphopeptides targeting the polo-box domain of polo-like kinase 1. Nat Struct Mol Biol. 2009;16:876–82.

35. Janeček M, Rossmann M, Sharma P, Emery A, Huggins DJ, Stockwell SR, et al. Allosteric modulation of AURKA kinase activity by a small-molecule inhibitor of its protein-protein interaction with TPX2. Sci Rep. 2016;6:28528.

36. Hu JH, Miller SM, Geurts MH, Tang W, Chen L, Sun N, et al. Evolved Cas9 variants with broad PAM compatibility and high DNA specificity. Nature. 2018;556:57–63.

37. Gaudelli NM, Komor AC, Rees HA, Packer MS, Badran AH, Bryson DI, et al. Programmable base editing of A•T to G•C in genomic DNA without DNA cleavage. Nature. 2017;551:464–71.

38. Komor AC, Kim YB, Packer MS, Zuris JA, Liu DR. Programmable editing of a target base in genomic DNA without double-stranded DNA cleavage. Nature. 2016;533:420–4.

39. R Development Core Team. R: A language and environment for statistical computing. R Foundation for Statistical Computing. 2008. http://www.r-project.org

40. Aken BL, Achuthan P, Akanni W, Amode MR, Bernsdorff F, Bhai J, et al. Ensembl 2017. Nucleic Acids Res. 2017;45:D635–42.

41. Love MI, Huber W, Anders S. Moderated estimation of fold change and dispersion for RNA-seq data with DESeq2. Genome Biol. 2014;15:550.

42. Morgens DW, Wainberg M, Boyle EA, Ursu O, Araya CL, Tsui CK, et al. Genome-scale measurement of off-target activity using Cas9 toxicity in high-throughput screens. Nat Commun. 2017;8:15178.

43. Hsu PD, Scott DA, Weinstein JA, Ran FA, Konermann S, Agarwala V, et al. DNA targeting specificity of RNA-guided Cas9 nucleases. Nat Biotechnol. 2013;31:827–32.

44. Rodriguez JM, Rodriguez-Rivas J, Di Domenico T, Vázquez J, Valencia A, Tress ML. APPRIS 2017: principal isoforms for multiple gene sets. Nucleic Acids Res. 2017.

45. He B, Wang K, Liu Y, Xue B, Uversky VN, Dunker AK. Predicting intrinsic disorder in proteins: an overview. Cell Res. 2009;19:929–49.

46. Cock PJA, Antao T, Chang JT, Chapman BA, Cox CJ, Dalke A, et al. Biopython: freely available Python tools for computational molecular biology and bioinformatics. Bioinformatics. 2009;25:1422–3.

47. Jones E, Oliphant E, Peterson P, et al. SciPy: Open Source Scientific Tools for Python. http://www.scipy.org/

48. Pedregosa F, Varoquaux G, Gramfort A, Michel V, Thirion B, Grisel O, et al. Scikit-learn: Machine Learning in Python. J Mach Learn Res. 2011;12:2825–30.

49. Ke G, Meng Q, Finley T, Wang T, Chen W, Ma W, et al. LightGBM: A Highly Efficient Gradient Boosting Decision Tree. Adv Neural Inf Process Syst. 2017;3149–57.

